# Crystallization and structure of ebselen bound to cysteine 141 of human inositol monophosphatase (IMPase)

**DOI:** 10.1101/2020.07.08.193284

**Authors:** Gareth D. Fenn, Helen Waller-Evans, John R. Atack, Benjamin D. Bax

## Abstract

Inositol monophosphatase (IMPase) is inhibited by lithium, the most efficacious treatment for bipolar disorder. Several therapies have been approved, or are going through clinical trials, aimed at the replacement of lithium in the treatment of bipolar disorder. One candidate small molecule is ebselen, a selenium-containing antioxidant, which has been demonstrated to produce lithium-like effects, both in a murine model and in clinical trials.

Here we present the crystallization and first structure of human IMPase covalently complexed with ebselen, a 1.47Å crystal structure (PDB entry 6ZK0). In the human-IMPase-complex ebselen, in a ring opened conformation, is covalently attached to Cys141, a residue located away from the active site.

IMPase is a dimeric enzyme and, in the crystal structure, two adjacent dimers share four ebselen molecules, creating a tetramer with ∼222 symmetry. In the crystal structure presented in this publication, the active site in the tetramer is still accessible, suggesting that ebselen may function as an allosteric inhibitor, or may block the binding of partner proteins.

**Synopsis:** Here we present a 1.47Å crystal structure of human inositol monophosphatase (IMPase) bound to the inhibitor ebselen (PDB entry 6ZK0). In the structure, ebselen forms a seleno-sulfide bond with cysteine 141 and ebselen-mediated contacts between two dimers give a ∼222 tetramer.

## 1. Introduction

Bipolar disorder is a chronic and debilitating psychiatric disorder, characterised by cycles of mania followed by severe depression, frequently accompanied by bouts of psychosis. Although antipsychotic agents are the preferred short-term method of treatment, more efficacious mood stabilising drugs, such as lithium, are used in long term clinical management (Geddes & Miklowitz, 2013). Lithium is the gold-standard treatment for bipolar, however, it has several serious side effects, such as nausea and cognitive impairment; in addition to a narrow therapeutic window (Rybakowski, 2016). Because of these liabilities, other, less efficacious mood stabilisers (e.g. Lamotrigine), are now often used in the treatment of bipolar disorder (Won & Kim, 2017). One enzyme inhibited by lithium, is inositol monophosphatase (IMPase) (Gill, *et al*. 2005), which has led to rational drug design targeting IMPase as a strategy for developing novel therapies for bipolar disorder (Brown & Tracy, 2013).

IMPase is a key enzyme in the phosphatidylinositol intracellular (PI) signalling pathway, whereby IMPase dephosphorylates inositol 1-, 3-, or 4-phosphate, collectively known as InsP1, to produce myo-inositol, also known as free inositol (Atack *et al*., 1995). Cleavage of InsP1 into myo-inositol by IMPase is required for the recycling of inositol for subsequent use in the PI signalling pathway (Atack *et al*., 1995). Inositol is an essential precursor for the synthesis of PI, subsequently utilised in the synthesis of phosphatidylinositol phosphates (PIPs). These include PI(4,5)P_2_, which is cleaved by phospholipase C following GPCR signalling to release the second messengers diacylglycerol (DAG) and inositol 1,4,5-trisphosphate (IP_3_) (Phiel & Klein, 2001). The observed depletion of free inositol and accumulation of the substrate of IMPase, InsP1, coupled with a reduction in agonist-invoked IP_3_ formation in cells and animals treated with lithium, led to the development of the inositol depletion hypothesis to explain the mechanism by which lithium exerts its effects (Berridge *et al*., 1989).

The inositol depletion hypothesis, suggests that lithium produces a reduction in free inositol, primarily via blocking the recycling of inositol from InsP1, which leads to a decrease in PI(4,5)P_2_ and a slowing of the PIP signalling pathways that are postulated to be hyperactive in bipolar disorder (Harwood, 2005). Further evidence to support the inositol depletion hypothesis come from observations that the mood stabilisers carbamazepine and valproic acid, also lead to depletions in free inositol and attenuation of PI(4,5)P_2_ signalling pathways (Williams *et al*., 2002). Therefore, targeting IMPase as a means of depleting free inositol has been of scientific interest, and has led to the search for new IMPase inhibitors.

One such IMPase inhibitor is ebselen (2-phenyl-1,2-benzisoselenazol-3(2H)-one), an organoselenium compound, which functions as a glutathione peroxidase mimic (Nakamura *et al*., 2002). Ebselen is believed to act through reduction of reactive oxygen species (ROS), by binding covalently to cysteine residues or thiols to form seleno-sulfide bonds that lead to the pharmacological effect (Azad & Tomar, 2014). However, it is not known whether the covalent binding of ebselen to specific groups is directly or indirectly responsible for its mechanism of action (Ulrich *et al*., 1996).

Ebselen has been demonstrated to inhibit IMPase in a covalent manner, with effects consistent with that of lithium, through depletion of free inositol in mouse brain (Singh *et al*., 2013). Subsequent trials in a healthy cohort demonstrated that ebselen leads to decreased myo-inositol in the anterior cingulate cortex, in addition to effects consistent with attenuation of PIP signalling (Singh *et al*., 2015). At present ebselen is currently in stage 2 clinical trial for the treatment of bipolar disorder, however, results from the trial have not been released at the time of writing.

Ebselen is known to bind to several proteins; crystal structures show ebselen covalently bound to cysteine residues of proteins including SOD1 (Capper., *et al* 2018; Chantadul *et al* 2020) and the transpeptidase LdtMt2 from *Mycobacterium tuberculosis* (De Munnik., *et al* 2019). Ebselen has also been reported to inhibit the Main protease from SARS-CoV-2 (Jin., *et al* 2020). These multiple targets suggest several potential therapeutic uses for ebselen, but also that there are likely to be off-target side effects.

Whilst inhibition of IMPase by ebselen *in-vitro* has been demonstrated, confirmation of ebselen binding loci and the exact mechanism of action on IMPase remains unclear. Structures of IMPase have been published with a variety of ligands, including a structure of human IMPase with the lithium mimetic L-690,330 (Kraft *et al*., 2018). In this paper, we present a 1.47Å structure of ebselen covalently bound to cysteine residue 141 of human IMPase (PDB entry 6ZK0). This is the first structure of IMPase to be published demonstrating direct covalent binding of ebselen to IMPase.

## 2. Materials and methods

All reagents were purchased from Sigma-Aldrich or ThermoFisher unless otherwise stated.

### 2.1. Macromolecule production

The IMPase construct was described by Kraft *et al*. 2018, with Rossetta2 (DE3) *E. coli* used for IMPase production. A starter culture (10 ml) of transformed *E. coli* was grown overnight and used to inoculate 1L of LB medium containing 2.5 mM betaine, 660 mM sorbitol, 35 mg/ml chloramphenicol and 50 mg/ml ampicillin. The culture was grown at 37 °C to an OD_600_ 0.8. IMPase expression was induced by the addition of 0.5 mM IPTG, and the culture grown overnight at 25 °C.

*E. coli* cells were pelleted by centrifugation at 9,600g for 20 min at 4 °C, and resuspended in 50 ml lysis buffer [20 mM Tris–HCl pH 7.8, 150 mM NaCl, 40 U/ml DNAse and one EDTA-free protease-inhibitor tablet (Roche)]. The resuspended pellet was lysed by sonication, and the debris pelleted by centrifugation at 33,000g for 20 min at 4 °C. The clarified sonicate was heat-treated at 68 °C for 1 h and the precipitant pelleted by centrifugation at 32,000g for 20 min at 4 °C. The supernatant was then incubated at 4 °C for 1 h with Co-NTA resin. After incubation, the resin was washed with seven resin volumes (RV) wash buffer [20 mM Tris–HCl pH 7.8, 150 mM NaCl, 15 mM imidazole] and IMPase eluted with 5 RV elution buffer [20 mM Tris–HCl pH 7.8, 150 mM NaCl, 250 mM imidazole].

The eluted fractions were pooled and incubated overnight at 4 °C with Pierce HRV 3C Protease solution. This removes the N-terminal tag (*MHHHHHHLEVLFQ*) by cleaving LEVLFQ↓GP (sequence in Table 1). Cleaved IMPase was purified initially by incubating with Glutathione Sepharose 4 Fast Flow Resin that had been pre-equilibrated with two column volumes of size-exclusion chromatography (SEC) buffer (20 mM Tris–HCl pH 7.8, 150 mM NaCl). The flow through from the column was collected and concentrated to 5 ml and loaded onto a HiLoad 26/600 Superdex 75 prep-grade SEC column pre-equilibrated with SEC buffer. Proteins were gel-filtered at a flow rate of 1 ml min^-1^ and the fractions containing IMPase were: pooled, buffer exchanged into storage buffer [20 mM Tris–HCl pH 7.8, 150 mM NaCl, 1 mM EDTA, 10%(v/v) glycerol] and concentrated to 20 mg/ml using 10 kDa protein concentrators at 4,000g, prior to storage at −20 °C. This protocol gave a typical yield of 2 mg IMPase per litre of culture.

**Table 1.**
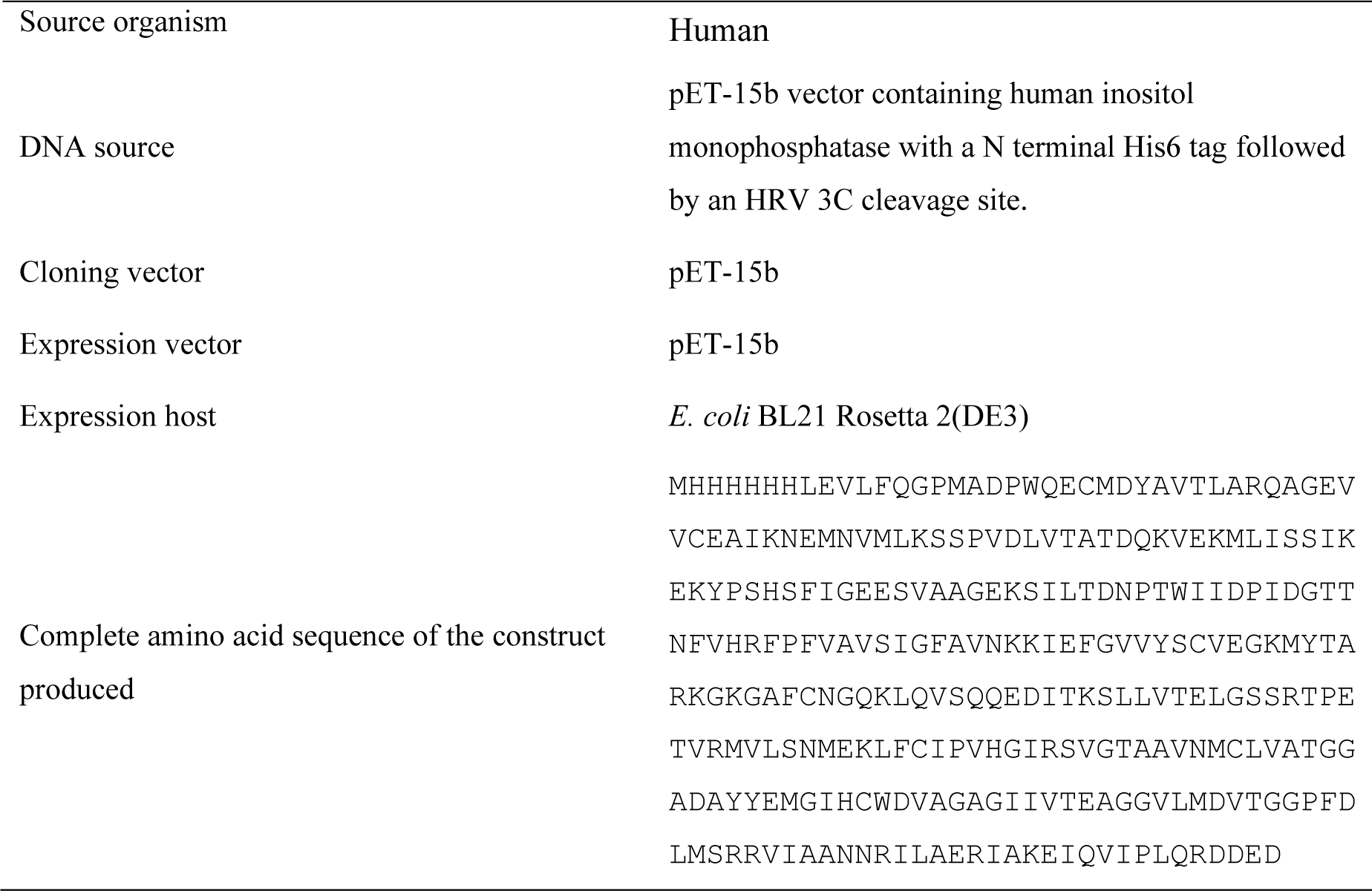
Macromolecule production information.

### 2.2. Crystallization

Human IMPase crystallisation was carried out using Swissci 3 lens sitting drop plates, and the plates set up with a mosquito liquid handling robot (TTP Labtech). To co-crystallise IMPase with ebselen, a 20 μl aliquot of IMPase, 20 mg/ml, in storage buffer [20 mM Tris–HCl pH 7.8, 150 mM NaCl, 1 mM EDTA, 10%(v/v) glycerol] was incubated with 50 mM ebselen (6 μl of 200 mM ebselen in DMSO). The sample was transferred to a 500 μl microcentrifuge tube and placed on a roller and incubated at room temperature for 30 min, prior to setting up crystallisation plates

A reservoir solution comprising the following: 0.2 M MnSO_4_, 0.1 M MES, 28% PEG4000, pH 5.5, was added to the IMPase and ebselen solution at a 1:1 ratio and incubated at 20 °C (see Table 2). Crystals appeared after 7 days and continued to grow until being harvested for data collection on day 14; being cryoprotected by transfer into 20% (v/v) glycerol + reservoir solution (80% v/v) and flash cooled prior to data collection.

**Table 2.** Crystallization conditions.

|  |  |
| --- | --- |
| Method | Sitting-drop vapour diffusion |
| Plate type | SWISSCI 3 lens crystallisation plate |
| Temperature (K) | 293 |
| Protein concentration | 20 mg/ml |
| Buffer composition of protein solution | 20 mM Tris–HCl pH 7.8, 150 mM NaCl, 1 mM EDTA, 10% (v/v) glycerol |
| Composition of reservoir solution | 0.2 M $\text{MnSO}_4$, 0.1 M MES, 28% PEG4000, pH 5.5 |
| Volume and ratio of drop | 100 nL, 1:1 |
| Volume of reservoir | 50 μL |

### 2.3. Data collection and processing

An X-ray diffraction dataset was collected from a single cryo-cooled crystal on beam-line I04-1 at the Diamond Light Source Synchrotron (Table 3). The data (2000, 0.1degree images) were processed with the Xia2 pipeline at Diamond (Winter, 2010) to give a 1.47 Å dataset in P 3_1_21 with cell dimensions a=b=84.02 Å, c=150.22 Å, α=β=90°, γ=120°.

**Table 3.**
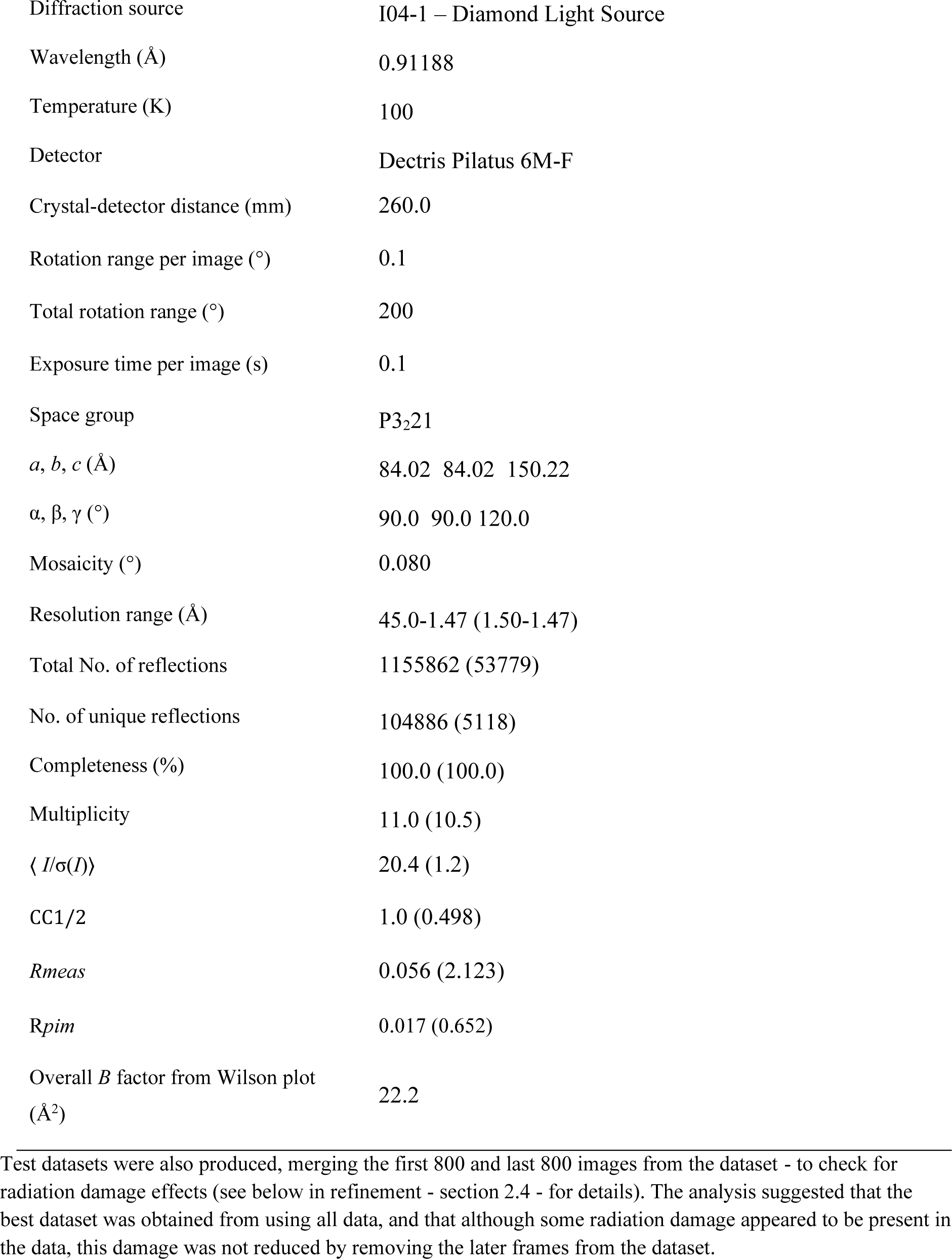
Data collection and processing. Values for the outer shell are given in parentheses.

**Table 4.** Structure solution and refinement. Values for the outer shell are given in parentheses.

|  |  |
| --- | --- |
| Resolution range (Å) | 24.44-1.47 (1.50-1.47) |
| Completeness (%) | 100.0 (100.0) |
| σ cutoff | NA |
| No. of reflections, working set | 99740 |
| No. of reflections, test set | 5081 |
| Final $R_{\text{cryst}}$ | 0.1734 |
| Final $R_{\text{free}}$ | 0.2011 |
| Cruickshank DPI | 0.0662 |
| No. of non-H atoms |  |
| Protein | 3694 |
| Ebselen* | 49 |
| Ligand | 54 |
| Water | 416 |
| R.m.s. deviations |  |
| Bonds (Å) | 0.014 |
| Angles (°) | 1.856 |
| Average $B$ factors (Å<sup>2</sup>) | 29.4 |
| Protein | 27.9 |
| Ebselen | 31.7 |
| Ligand | 45.7 |
| Water | 40.3 |
| Ramachandran plot |  |
| Favoured regions (%) | 98 |
| Additionally allowed (%) | 1.5 |
| Outliers(%)** | 0.5 |

\* Ebselen - attached to Cys141A had only one atom with two positions (the selenium atom). Ebselen attached to Cys141B had every atom in two positions. There are sixteen non-hydrogen atoms in ebselen.
\*\* Lys36 is the only residue (just) outside allowed region in both subunits ( $\phi/\psi$ -100°/-110°)

There are nine crystal structures of human IMPase in the PDB with similar cell dimensions, all in space-group P3_2_21; thus, the data were reindexed and remerged in P3_2_21. The data were merged with version 0.7.4 of the program AIMLESS (Evans & Murshudov, 2013). The Rcp statistic, used to estimate cumulative radiation damage in AIMLESS (Diederichs, 2005), did not increase significantly over the 2000 frames.

### 2.4. Structure solution and refinement

The structure was solved by rigid body refinement from the 1.7 Å crystal structure of IMPase with Mn (PDB entry 6GJ0; Kraft *et al*., 2018). Initial structure solution used the DIMPLE pipeline; this pipeline provides the user with a quick method to identify datasets that have a bound ligand or drug candidate in their crystal (http://ccp4.github.io/dimple/; Wojdyr *et al*., 2013).

Initial maps showed clear electron density (Figure 1) for a single ebselen molecule attached to Cys141 in both subunits A and B of the dimer (the P3_2_21 cell has one dimer in the asymmetric unit). The ebselen was built onto Cys141A and Cys141B in coot (Emsley *et al*., 2010; Emsley, 2017). Restraints for the covalently bound ebselen were generated in AceDRG (Long *et al*., 2017) and the structure was refined using Refmac5 (Murshudov *et al*., 2011).

**Figure 1.**
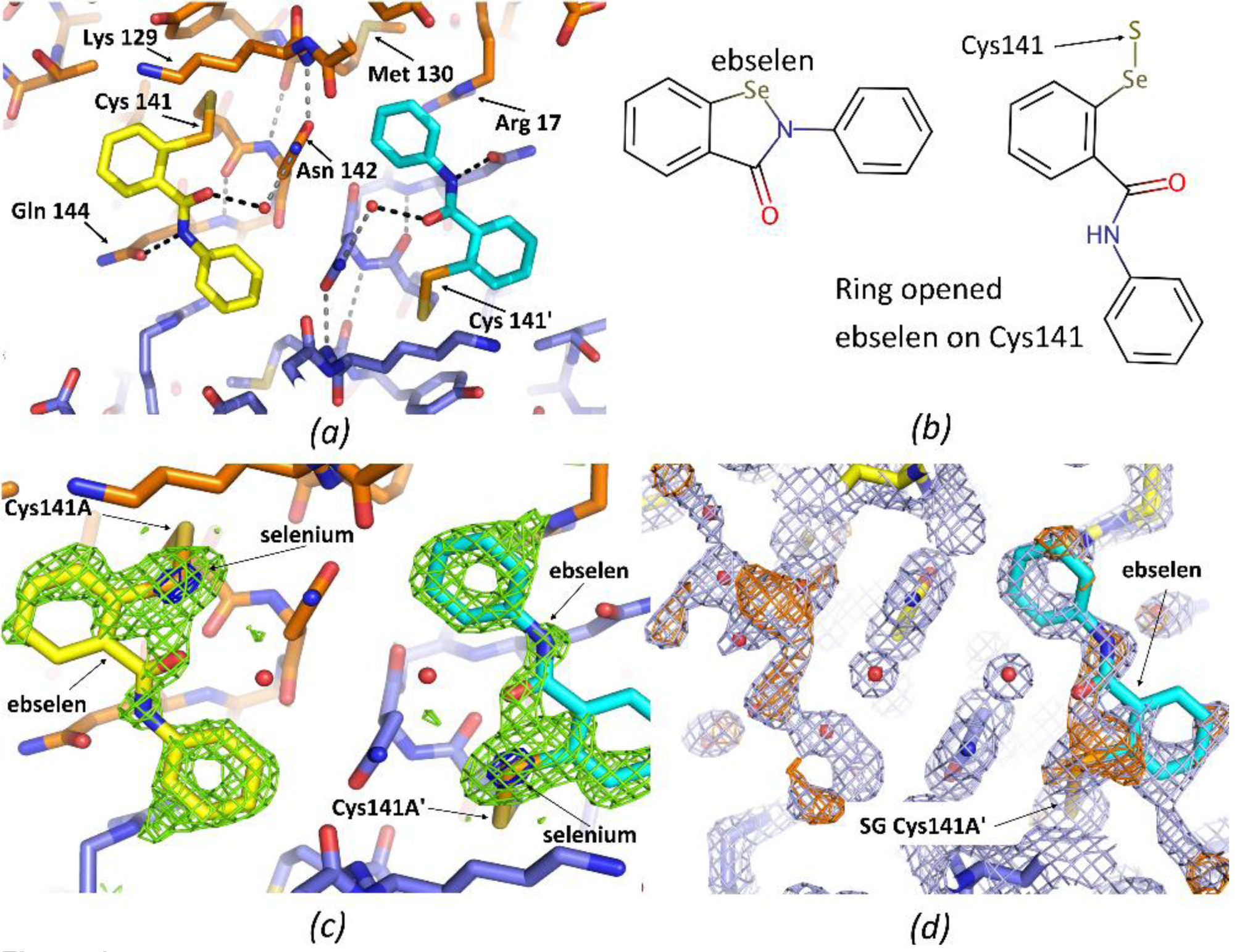
View of ebselen attached to Cys141 (PDB entry 6ZK0). *(a)* Overview of two ebselen molecules attached to A and A’ (symmetry related) subunits around crystallographic twofold axis. One subunit (A) has carbons in orange and the second (A’) subunit has carbons slate-blue (nitrogens are blue, carbons red, seleniums orange and sulphurs yellow-orange). Carbons in one ebselen are yellow and are cyan in the second ebselen. Hydrogen bonds near ebselen are indicated by dotted lines. *(b)* chemical structures of ebselen and ring-open ebselen on Cys141(drawn with Marvin, https://www.chemaxon.com). *(c)* Final ebselen omit map (Fo-Fc) (3 sigma green, 15 sigma blue). Note peaks on seleniums (blue mesh) are 20.5 and 19.6 sigma in this ebselen omit map. *(d* Original DIMPLE (Wojdyr *et al*., 2013) 2Fo-Fc map (1sigma – light blue), and difference map Fo-Fc (3 sigma - orange). For subunit A the DIMPLE refined structure with waters (small red spheres) refined into the density for the ebselen is shown. For the A’ subunit the ‘final’ coordinates (including ebselen) are shown.

Although the electron density was very clear for the terminal phenyl ring of the compound (Figure 1c and 1d) and electron density maps have a large peak for the selenium (covalently bonded to the sulfur of Cys141), the electron density suggests some radiation damage has occurred to the sulfur-selenium bond (Weik *et al*., 2002; Garman, 2010). The data are consistent with a model in which some initial radiation damage occurred to the sulfur-selenium (within the first few degrees), after which a steady state occurred (bond reforming after breaking due to radiation) (Gerstel *et al*., 2015).

Each active site contains three Mn^2+^ ions in 6GJ0 (Kraft *et al*., 2018), while in 2BJI (Gill *et al*., 2005) each active site contains three Mg^2+^ ions. In our structure, site 2 (Gill *et al*., 2005) does not have enough electron density for a Mn^2+^ ion (Mn^2+^ ions have twenty-three electrons). We modelled a similarly coordinated sodium ion at this site, because we have no Mg^2+^ ions in our crystallisation experiment (the protein comes in 150mM NaCl and the crystallisation buffer contains 200mM MnSO_4_). However, we cannot rule out the possibility that this is a Mg^2+^ ion, rather than a Na^+^ ion (both Na^+^ and Mg^2+^ ions have ten electrons). This ‘site 2’ is the position where lithium is postulated to bind with tetrahedral coordination geometry (Gill *et al*., 2005). However, the coordination geometry in our structure is consistent with two Mn^2+^ ions and one Mg^2+^ ion each with ‘standard’ octahedral coordination geometry. Most of the active site metal atoms are modelled in two positions and have temperature factors similar to those of surrounding residues (Masmaliyeva & Murshudov, 2019).

In the deposited structure (PDB 6ZK0), the ebselens on Cys141 in subunits A and B each have an occupancy of 0.6, but the selenium atoms are modelled in two positions (supplementary Figure 1). In the crystal, the selenium-sulfur bond has been modelled with an occupancy of 0.4 or 0.35 (see supplementary Figure 1). A second selenium position, observed in electron density maps, is some 1.3 Å further away from the sulfur of Cys141 and this second position is likely caused by radiation induced cleavage of the seleno-sulfide bond (Gerstel *et al*., 2015).

## 3. Results

### 3.1. Human IMPase structure with ebselen on cysteine 141

The 1.47 Å crystal structure of human IMPase with ebselen (PDB entry 6ZK0) was solved from a structure of human IMPase in the same unit cell and space-group (6GJ0 - Kraft *et al*., 2018). Electron density maps (Figure 1) clearly showed ebselen attached only to a single cysteine, Cys141. The binding of ebselen to the Cys141 residue in each monomer does not lead to noticeable changes in conformation in the active site that would prevent the catalytic activity of IMPase. Additionally, the binding of ebselen to Cys141 does not appear to prevent dimer formation, as evidenced by the dimers (and tetramers) present in this structure (6ZK0). However, whilst this structure clearly shows ebselen bound to Cys141 on each monomer of IMPase, our structure does not rule out the possibility that Cys218 could be modified *in-vivo*.

IMPase contains seven cysteine residues, amino acids 8, 24, 125, 141, 184, 201 and 218, of these residues, only Cys218 is near the active site. Four of the cysteine residues are buried and would not be expected to be accessible to modification by ebselen (cysteines 8, 125, 201 and 218). Of the three cysteines that have some surface accessibility in the monomer, one of them, Cys184 is largely buried in the dimer interface as shown in Figure 2 (PDB entry 6ZK0). The procedure used to co-crystallise ebselen with IMPase allowed partial oxidation to form the Cys141 - ebselen seleno-sulfide bond (Fig 1.). In our structure Cys24 has some surface accessibility when reduced, but when oxidised forms a disulphide with Cys125 (Figure 2). Cys125 has no surface accessibility whether oxidised or reduced. A partial Cys25-Cys125 disulphide is also observed in 6GJ0 (Kraft *et al*., 2018).

**Figure 2.**
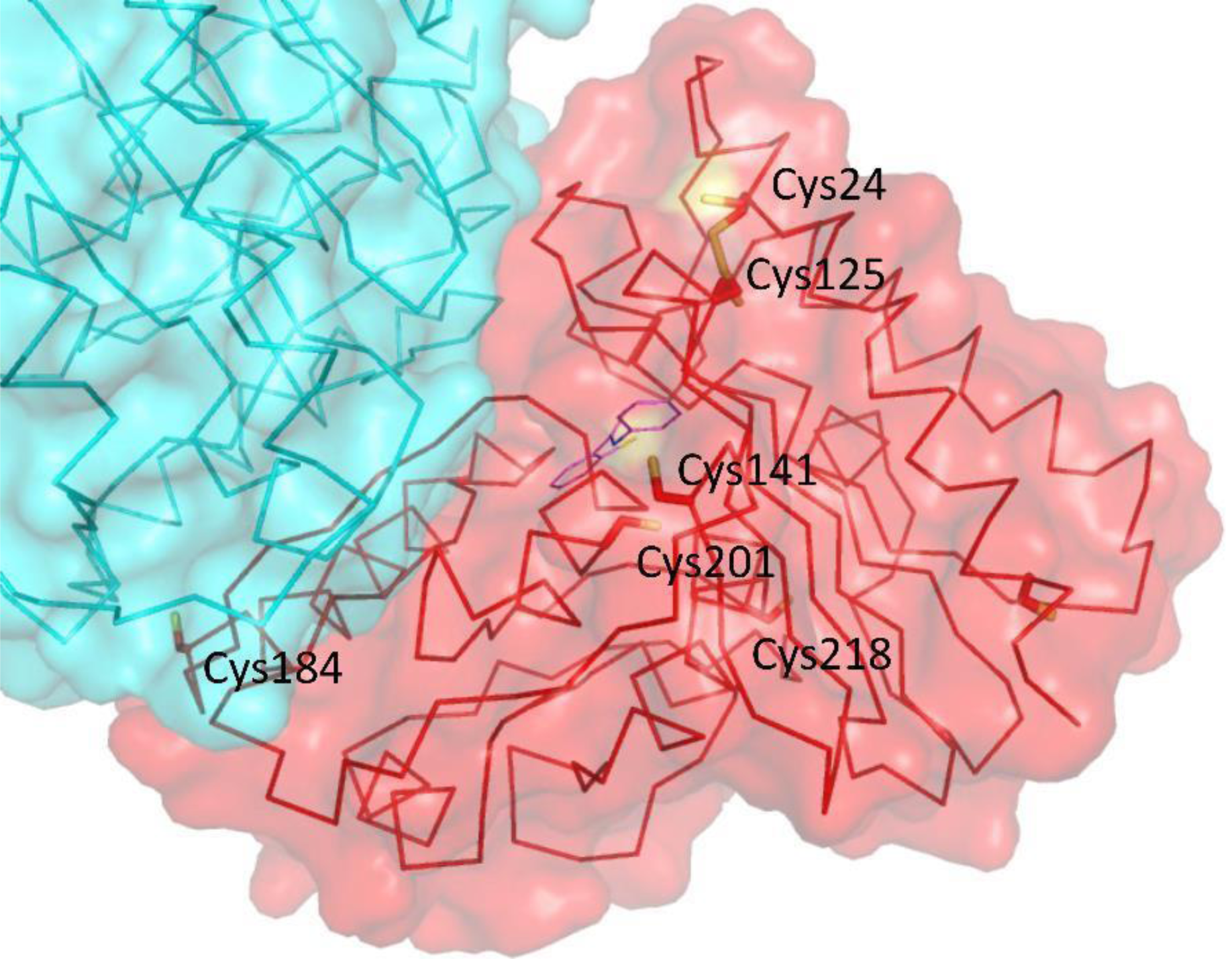
Cysteine residues in IMPase with ebselen bound to Cys141 (PDB entry 6ZK0). IMPase is shown as with a Ca ribbon trace and the side-chains of the seven cysteines are shown in stick on the ‘red’ subunit. The second subunit in the dimer is shown in cyan. A semi-transparent surface is shown; note that where the sulphurs of the cysteine residues are on the surface of the protein, the surface is yellow (Cys24 and Cys141). Cys184 also has some surface accessibility in the monomer, but is largely buried at the dimer interface, so no yellow is visible for Cys184 in this figure.

Previous research suggested that Cys218 is the primary reactive cysteine residue (Knowles *et al*., 1992), supported by reduced inhibition of C218A IMPase by ebselen (Singh *et al*., 2013). In the structure reported here (PDB entry 6ZK0), and other human IMPase structures, Cys218 is largely buried, and therefore seems an unlikely target for modification, as it is unclear as to how ebselen would gain access. If the side-chain of Cys218 is modified by ebselen it would likely lead to substantial reorganisation of the protein structure; Asp220 coordinates an active site metal ion.

From our structure, it appears that Cys141 is likely to be the primary binding site of ebselen. The sulfur group of this residue is exposed on the surface of IMPase (Figure 2), and this residue is not in close proximity to another cysteine residue, in either the monomeric or dimer form, so unlikely to form a disulphide bond (PDB entry 6ZK0). Cys141 is conserved in mammals (Knowles *et al*., 1992; Singh *et al*., 2013), and an analogous cysteine residue (Cys138) is present in *Staphylococcus aureus* IMPase. Given this level of conservation, it is probable that Cys141 is a functionally important residue in IMPase (Dutta *et al*., 2014).

Cys141 has previously been shown to be a reactive cysteine residue, demonstrated through affinity for the thiol probes pyrene-maleimide (Greasley *et al*., 1994) and n-ethylmaleimide (Knowles *et al*., 1992). In this structure, ebselen is in an open ring conformation with the selenium atom forming a seleno-sulfide bond with the sulfur group of Cys141, which is consistent with the known binding mechanism of ebselen (Capper *et al*., 2018). Each monomer of IMPase in this structure has a single ebselen molecule bound to Cys141 (PDB entry 6ZK0).

Whilst Cys141 conservation would suggest a critical role of this residue, the exact function remains unclear. The residue is not in close proximity to the active site, therefore the residue is unlikely to be involved in catalytic activity. One possibility is that the residue may undergo post translational modification *in-vivo*, and may be redox active (Marino & Gladyshev, 2012), linking ebselen to therapeutic effects as a known antioxidant.

### 3.2. Ebselen bound IMPase is observed to form a tetramer with 222 symmetry in the solid state

Figure 3 shows the position of Cys141 on the subunits A and B in an IMPase dimer. The two ebselen molecules covalently bound to each dimer and localise with two other ebselen molecules on a second dimer. The 2 ebselen molecules on each dimer increase contacts with a neighbouring dimer, which gives a tetramer with ∼222 symmetry in the crystal. However, as shown in Figure 3 (panels g and h), the active site still appears accessible.

**Figure 3.**
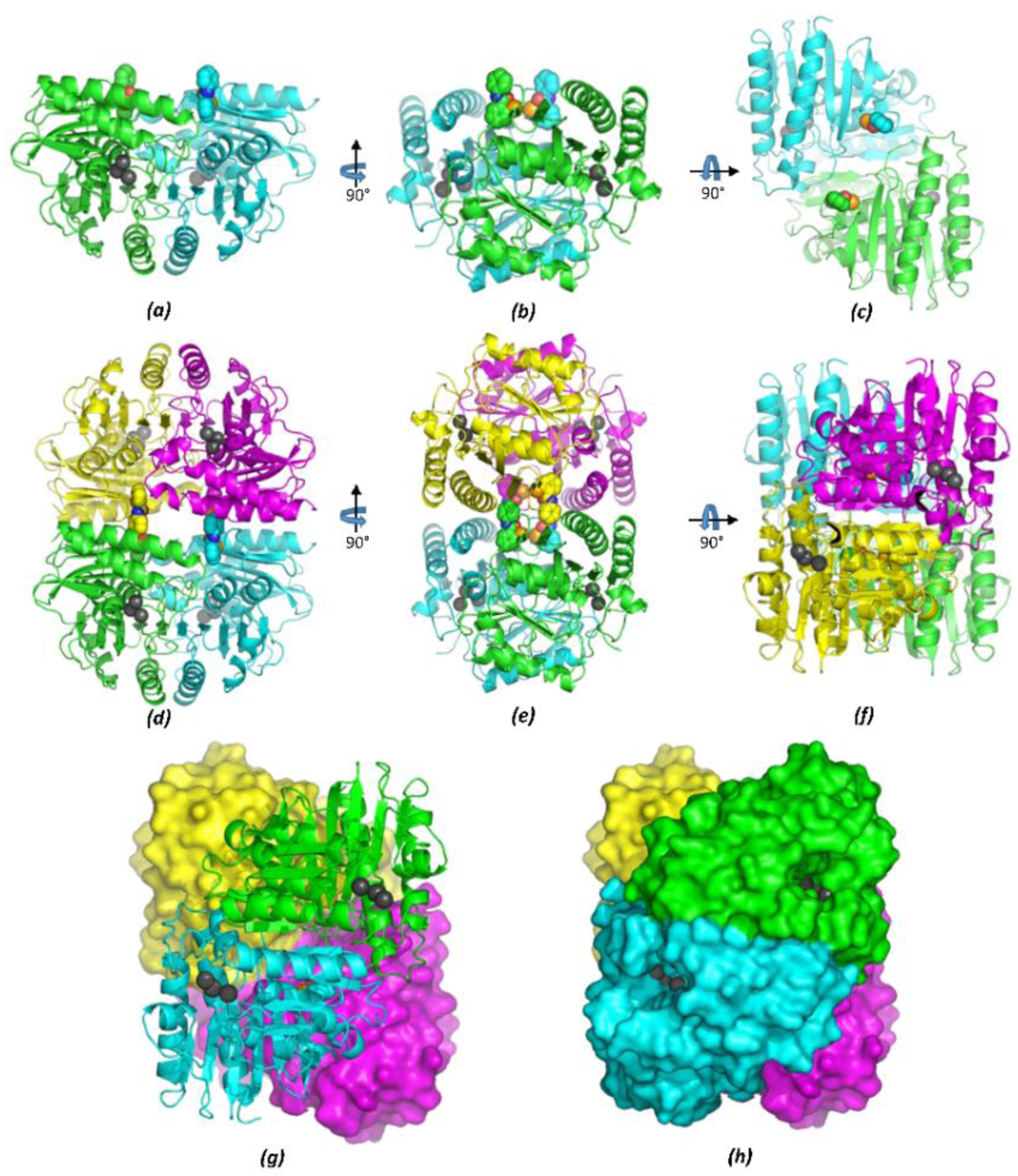
Orthogonal views of IMPase dimer (and tetramer) showing ebselen on Cys141 and metal ions in the active sites based on the structure of PDB entry 6ZK0. *(a)* The two subunits in the dimer are shown as green (subunit A) and cyan (subunit B) cartoons with ebselen attached to Cys141A and Cys141B in space-fill. Metal ions (Mn^2+^/Na^+^) at each active site are shown as grey spheres. *(b)* and *(c)* - orthogonal views. *(d),(e),(f)*. Same views as *(a),(b),(c)* but also showing a second dimer related by a crystallographic twofold axis. Subunit A’ is in yellow and B’ in magenta. The view in *(e)* is along the crystallographic twofold that rotates the A-B dimer (green/cyan) onto the A’-B’ dimer (yellow-magenta). *(g), (h)*. Two views of the tetramer from ‘underneath’. Showing that the three metal ions (grey/black spheres) at each active site are still accessible in the tetramer. In view *(h)*, a surface is shown on both dimers.

Interestingly, other members of the IMPase superfamily, including Fructose-1,6-bisphosphate (FBPase), have been observed to have both dimer and tetramer forms (Hines *et al*., 2007). Tetrameric forms of IMPase have also been observed in the anaerobic hyperthermophilic eubacterium *T. maritima* (Stieglitz *et al*., 2007).

## 4. Discussion

The IMPase crystal structure that is presented (PDB entry 6ZK0) has ebselen covalently attached to Cys141, however it is not clear to what extent this binding brings about ebselen’s inhibitory effects on IMPase. It is possible that the modification on Cys141 is biologically relevant, and that this is a cysteine residue that is modified *in vivo* by ebselen, with binding over the previously suggested preferred residue Cys218 (Singh *et al*., 2013). Cys141 has been shown to be a reactive cysteine residue (Greasley *et al*., 1994), so it possible that ebselen binding at Cys141 causes inhibition of IMPase. However, the binding of ebselen does not affect the conformation of the active site or prevent dimer formation; with the high conservation of Cys141 across species, it is likely to be a functional residue with a potential regulatory redox role.

There is evidence that modulation of IMPase away from the active site and dimer interface can affect activity; synthetic peptides that disrupt IMPase-calbindin interactions prevent calbindin mediated activation of IMPase (Noble et al., 2018) and mediate antidepressant-like effects in mice (Levi *et al*. 2013). It is possible that ebselen interferes with accessory protein binding, possibly by formation of the tetramer seen in the crystal, to moderate the activity of IMPase *in-vivo*.

However, the possibility that the conditions used in crystallisation do not reflect physiological conditions and that, *in vivo*, Cys141 and Cys218 could be modified differently by ebselen, cannot be ruled out. The binding of ebselen to Cys141 does not appear to have significantly altered the structure of the dimer or the active site. Should the ebselen/IMPase tetramers observed prove to be biologically relevant, this suggests a new mechanism for the regulation and subsequent inhibition of IMPase that could be utilised in the development of novel therapeutics.

## Acknowledgements

We are especially grateful to Dr. Pierre Rizkullah and his colleagues for transporting and carrying out data collection on our crystals. We also thank Diamond Light Source Ltd (Didcot, UK) for access to synchrotron radiation on beamline I04. We thank Gareth Wright for helpful discussions.

